# Wild-type IDH1 inhibition enhances chemotherapy response in melanoma

**DOI:** 10.1101/2022.05.13.491880

**Authors:** Mehrdad Zarei, Omid Hajihassani, Jonathan J. Hue, Hallie J. Graor, Moeez Rathore, Ali Vaziri-Gohar, John M. Asara, Jordan M. Winter, Luke D. Rothermel

## Abstract

Malignant melanoma is one of the most common types of cancer in the United States. Despite recent and well-described progress in melanoma treatment, advanced disease still carries a poor prognosis for many patients and chemotherapy has been appropriately abandoned as a front-line option. Wild-type isocitrate dehydrogenase 1 (wtIDH1) has recently been implicated as a metabolic dependency in cancer. The enzyme is protective to cancer cells under metabolic stress, including oxidative damage by conventional chemotherapy and nutrient limitation characteristic of the tumor microenvironment. Specifically, the cytosolic enzyme generates NADPH to maintain redox homeostasis. IDH1 also supports mitochondrial function through anaplerosis of its reaction product, α-ketoglutarate. We show that melanoma patients express higher levels of the wtIDH1 enzyme compared to normal skin tissue, and elevated wtIDH1 expression portends poor patient survival. Knockdown of IDH1 by RNA interference inhibited cell proliferation and migration under low nutrient levels. Suppression of IDH1 expression in melanoma also decreased NADPH and glutathione levels, resulting in increased reactive oxygen species. An FDA-approved inhibitor of mutant IDH1, ivosidenib (AG-120), exhibited potent anti-wtIDH1 properties under low magnesium and nutrient levels, reflective of the tumor microenvironment *in natura*. Similarly, findings were replicated in murine models of melanoma. Further, wtIDH1 inhibition was synergistic to conventional anti-melanoma chemotherapy in pre-clinical models. This work points to a novel and readily available combination treatment strategy for patients with advanced and refractory melanoma.

## Introduction

Prior to the use of contemporary immunotherapy and targeted therapies, treatment options for advanced melanoma were largely restricted to conventional chemotherapeutics. Dacarbazine (DTIC), an FDA-approved agent in melanoma, produced an objective response rate of 13% to 20%, with a median survival rate of just 5 to 6 months for patients with stage IV disease (1). A DTIC derivative, temozolomide (TMZ), was comparable with respect to efficacy, but carried advantages of oral delivery and the ability to penetrate the blood-brain barrier (2). Both are prodrugs of the active alkylating agent 5-(3-methyltriazen-1-yl) imidazole-4-carboxamide (MTIC) that induces apoptosis through direct DNA damage (3).

Due to the availability of more effective therapeutics in the modern treatment era, chemotherapies are currently reserved for unique treatment scenarios (4,5). The five-year survival rate among patients with metastatic melanoma receiving combination immunotherapy with nivolumab and ipilimumab is over 50%, as compared to just 5% prior to the adoption of these therapies (6–8). Despite clear progress, 40% of patients treated with dual checkpoint inhibitor therapy do not experience any response, and 60% of patients experience significant toxicities from therapy. In addition, over one-third of responders (20% overall) eventually develop secondary or acquired resistance (9–11). Targeted therapies, such as BRAF and MEK inhibitors, have demonstrated success in patients with BRAF mutated tumors. However, only half of melanoma patients carry this mutation (12). In such patients, the overall survival benefits of these targeted therapies are modest, and acquired resistance occurs for almost all patients during the first year of treatment (13). Thus, while novel therapies have legitimately improved survival for patients with advanced melanoma, innate and acquired treatment resistance limits their generalizability and effectiveness. Attention in the field has concentrated heavily on these two areas of focus over the past decade (immunotherapy and oncogene-targeted therapy) at the expense of investigating alternative strategies to exploit critical biologic dependencies.

For instance, the harsh tumor microenvironment in melanoma is characterized by hypoxia, tissue necrosis, and nutrient limitation. These conditions require adaptive metabolic reprogramming for cancer cell survival (14–17). Among other biologic processes, cancer cells rely on robust antioxidant defense to neutralize reactive oxygen species generated under severely austere conditions, as well as enhanced mitochondria function to maximize ATP production in the face of nutrient scarcity (18,19). Recently, our group identified wild-type isocitrate dehydrogenase 1 (wtIDH1) as a key metabolic enzyme for both of these pro-survival cellular activities in pancreatic cancer (20). IDH1 is cytosolic and isofunctional to the mitochondrial enzymes IDH2 and IDH3A. The enzyme catalyzes the interconversion of isocitrate and alpha-ketoglutarate (αKG) using NADP(H) as a cofactor (21–23). Under nutrient limitation, commonly present in tumors such as melanoma, oxidative decarboxylation of isocitrate is favored, producing NADPH and αKG. These products directly support antioxidant defense (NADPH is the reductive currency in cells) and mitochondrial function (αKG fuels the TCA cycle through anaplerosis), respectively (24).

Our recent studies in pancreatic cancer identified for the first time that small molecules developed as selective mutant-IDH1 inhibitors (25–28), actually inhibit wtIDH1 with a high degree of potency under conditions present in the tumor microenvironment (24). Specifically, reduced magnesium levels in tumors allow mutant-IDH1 inhibitors to bind to the wtIDH1 allosteric site with greater affinity (29). In the presence of cancer-associated stress (e.g., nutrient limitation), cancer cells are highly dependent on wtIDH1, rendering wtIDH1 inhibition with allosteric IDH1 inhibitors lethal to treated cancer cells.

Previous work demonstrates that the induction of high levels of oxidative stress in melanoma cells can be exploited to overcome chemotherapy resistance, as antioxidant capabilities are overwhelmed (30). This paper is the first since our publication on pancreatic cancer (24) to validate the effectiveness of IDH1 inhibition in another cancer type. We build on upon this work to show that wtIDH1 is especially important for melanoma cell survival, and in particular, chemotherapy resistance. If true, this work provides a strong rationale to translate findings to clinical trials that test the combination of available wtIDH1 inhibitors with conventional chemotherapeutics largely abandoned for patients with advanced melanoma (e.g., DTIC or TMZ).

## Materials and Methods

### Cell lines and cell culture

A375 (human), SK-MEL-28 (human), and B16-F10 (murine) melanoma cell lines were obtained from ATCC (American Type Culture Collection). Cells were cultured in DMEM supplemented with 10% FBS (Gibco/Invitrogen), and 1% penicillin-streptomycin (Invitrogen) at 37°C in 5% humidified CO_2_ incubators. Glucose-free DMEM (Life Technologies, 21013-024) was utilized for experiments with varying glucose concentrations, and the appropriate amounts of glucose were added to the media. For experiments with varied magnesium levels, magnesium sulfate-depleted DMEM (Cell Culture Technologies, 964DME-0619) was utilized, and supplemented with the indicated amounts of MgSO4. Cell lines were treated with prophylactic doses of Plasmocin and Mycoplasma tested (# MP0035, Sigma Aldrich) monthly. Cell lines were passaged at least twice before experimental use.

### CRISPR construct knockout IDH1 in Melanoma cells

CRISPR/Cas9-mediated knockout of *IDH1* was performed in A375, and SK-MEL-28 cells using guide RNAs targeting *IDH1* (GTAGATCCAATTCCACGTAGGG) fused with CRISPR/Cas9 and GFP protein. CRISPR Universal Negative Control plasmid (CRISPR06-1EA) was purchased from Sigma-Aldrich (St. Louis, MO). Cells were collected after 48 hours of transfection, and GFP-positive cells were single-sorted using FACS ARIA flow cytometer.

### siRNA transfections

Cells were plated at 60% confluence in 6-well plates, and transient siRNA transfections (1μM) were performed using Lipofectamine 2000 (Invitrogen) and Opti-MEM (Invitrogen) according to the manufacturer’s protocol. Experiments were generally started 48 hours after transfections. Small interfering RNA (siRNA) oligos were purchased from Ambion (si*IDH1*, S7121; siCTRL, AM4635).

### Cell viability assays

Cells were seeded in 96-well plates with 1×10^3^ cells per well. After settling for 24 hours, cells were treated as indicated. Experiments lasted for 6 days unless otherwise detailed, and cell proliferation was estimated by staining with Quant-iT PicoGreen™ (Invitrogen). To estimate cell death, cells were trypsinized, stained with 0.4% Trypan blue (Invitrogen) after 0 to 4 days, and counted using a Hausser Scientific bright-line hemocytometer (Fisher Scientific).

Drug combination assays were performed after seeding 1-2×10^3^ cells per well in 96-well plates for 24 hours. Cells were treated with AG-120 (dose range: 0.125 μmol/ml - 2 μmol/ml) and TMZ (dose range: 6.25 μmol/ml - 800 μmol/ml) in a 6 × 8 well matrix, and experiments were repeated in triplicate. Cell viability was estimated after 6 days (compared to vehicle) with Quant-iT PicoGreen. Drug interactions were quantified and characterized as synergistic, additive, or antagonistic using the Bliss Independence model, as described (31). For all *in vitro* experiments using AG-120, cells were cultured under low magnesium conditions (<0.4 mM Mg^2+^) to effectively inhibit wtIDH1 enzyme activity (as a reference, normal culture media and serum contain roughly 1 mM Mg^2+^). Low glucose (2.5 mM glucose) was utilized as indicated to generate conditions of wtIDH1 dependency and simulate glucose levels in the tumor microenvironment (32–36).

### Immunoblotting

Cells were lysed using 1X RIPA buffer containing protease and phosphatase inhibitors. Protein concentration was quantified using the BCA Protein Assay (Thermo Fischer Scientific). Equal amounts of total protein were added to a 4–12% Bis-Tris gel (Life Technologies), separated by size using electrophoresis, and transferred to a PVDF membrane. Blots were blocked in 5% skimmed milk and probed with primary antibodies against anti-*IDH1* (Invitrogen, OTI2H9) and anti-α-tubulin (Invitrogen, 11224-1-AP). Chemiluminescent (32106, Thermo Fisher Scientific) signal was captured using a digital imager (Odyssey Imaging system).

### DNA sequencing

DNA extracted from 1×10^6^ human and murine melanoma cells using DNeasy Blood and Tissue kit (Qiagen) following the manufacturer’s protocol. A portion of IDH1 gene exon 4 containing Arg132 was amplified using set pairs of primers, against the human sequence: IDH1 F:5’-ACCAAATGGCACCATACGA-3; IDH1 R: 5’-TTCATACCTTGCTTAATGGGTGT-3’, and for mouse: IDH1 F:5’-ATTCTGGGTGGCACTGTCTT-3’; IDH1R: 5’-CTCTCTTAAGGGTGTAGATGCC-3’. PCR was performed using a DNA thermal cycler, and the products were analyzed by agarose gel electrophoresis. PCR products were sequenced using one of the amplification primers.

### Migration assay

Cells were plated at a density of 6×10^4^ cells in the upper chamber of a 6.5-mm Transwell with 8.0 μm pore polycarbonate membrane inserts (Corning). 100 μL of serum-free DMEM was added to the Transwells for 8 hours at 37 °C. Complete growth medium was placed in the bottom section as a chemoattractant. Non-migrated cells were wiped off the upper surface using cotton swabs. Cells migrating to the lower surface were fixed and stained using 0.5% crystal violet, imaged using a 10X objective on a Nikon TE200 microscope, and quantified using Image J analysis software.

### Clonogenic assay

Cells (2-3×10^3^ cells per well) were seeded in 6-well plates and treated with AG-120 (or vehicle) at the indicated concentration, and under low MgSO4 (0.08 mM) conditions. After 8 days, cells were washed with 1X PBS, fixed in 80% methanol, and stained with 0.03% (w/v) crystal violet for 10 minutes. The dye was extracted with 10% glacial acetic acid and absorbance was measured at 600 nm using a GloMax plate reader (Promega) (37).

### Cellular ROS and 8-OHdG analysis

Cells were seeded in 96-well black plates and incubated in 100 μL phenol red free media containing 10 μM H2-DCFDA (Invitrogen) for 45 min, at 37 °C, in the dark. Fluorescence was measured using an excitation wavelength at 485 nm and emission wavelength at 535 nm on a GloMax plate reader. 8-hydroxy-2-deoxyguanosine (8-OHdG) was measured (Abcam, AB201734) per the manufacturer’s instructions.

### Animal studies

All experiments involving mice were approved by the CWRU Institutional Animal Care Regulations and Use Committee (IACUC, protocol 2018-0063). Six-week-old female athymic nude mice (Nude-Foxn1nu) were purchased from Harlan Laboratories (6903M). A375 cells, or genetically modified variants, were suspended in 150 μL solution comprised of 60% Dulbecco’s PBS and 40% Matrigel. Suspensions of 1×10^6^ cells were injected subcutaneously into the right flank of mice. For syngeneic orthotopic experiments, 5×10^4^ B16-F10 cells were suspended in the same manner and injected into flanks of immunocompetent 10 week-old C57BL/6J mice.

Treatments were initiated after tumors were first palpable and reached 100-120 mm^3^ (nude mice) or 80-100 mm3 (C57BL/6J mice). AG-120 (Asta Tech, 40817) was administered orally at 150 mg/kg twice per day as a suspension in PEG-400, Tween-80, and saline (10:4:86). TMZ (Sigma-Aldrich, T2577) was given at 30 mg/kg as intraperitoneal injections, five times per week. Bodyweights and tumor volumes were measured weekly. For the latter, Vernier calipers were utilized and volumes estimated by the formula, Volume = (Length × Width^2^)/2. At the end of the experiment, mice were euthanized by carbon dioxide inhalation and tumors were immediately resected for additional studies. For immunohistochemistry analysis, tumors were fixed in 10% formalin (Thermo Fisher Scientific; 427-098) and stored at -80°C.

### Real-time quantitative PCR

RNA was extracted using the RNeasy PureLink RNA isolation (Life Technologies; 12183025) and converted to cDNA using a High-Capacity cDNA Reverse Transcription Kit, per the manufacturer’s protocol (Applied Biosystems; 4387406). qPCR was performed using Taqman™ Universal Master Mix II (Thermo Fisher Scientific; 4440038) with an IDH1 probe (Thermo Fisher Scientific; 4351372) and analyzed using the Bio-Rad CFX Maestro manager 2.0 software.

### Metabolites extraction and measurement by LCMS

Cells were grown to ∼50% confluence in complete growth medium in 6-well plates and in biological triplicates. After rinses with ice-cold PBS, metabolites were extracted with 80% HPLC-grade methanol, scraped, and collected. Polar metabolites were analyzed by 5500 QTRAP triple quadrupole mass spectrometry (AB/SCIEX) coupled to a Prominence UFLC HPLC system (Shimadzu) using amide HILIC chromatography (Waters) at pH 9.2, as previously described (38). 299 endogenous water-soluble metabolites were measured at a steady state. Data were normalized to protein content. NADPH (NADP/NADPH-Glo™ Promega G9081) and GSH levels (GSH-Glo^™^ Promega V6911) were also measured separately per the manufacturer’s instructions.

### Bioenergetics

Oxygen consumption rates (OCR) and extracellular acidification rates (ECAR) were quantified using the XFp mini extracellular analyzer (Seahorse Bioscience). A375 cells were seeded at 1×10^4^ cells per well in complete DMEM (25 mM glucose and 2 mM glutamine) in an Agilent XFp Cell Culture miniplate (#103025-100), and cultured at 37°C in 5% humidified CO_2_ incubators. For these experiments, glucose-free DMEM was supplemented with glucose to achieve the indicated concentrations, and incubated for an additional 36 hours. The XFp FluxPak cartridge (#<u>103022-100</u>), was hydrated and incubated at 37 °C, using non-CO_2_ incubator overnight. The following day, cells were washed twice and replaced with Seahorse XF base media (using the indicated glucose concentrations), and incubated in a non-CO_2_ incubator at 37°C. OCR and ECAR were measured in the standard fashion using standard mitochondrial inhibitors: 1.5 μM oligomycin, 2 μM FCCP, and 0.5 μM rotenone + 0.5 μM antimycin A (Mito Stress Test, #103015-100). Data were normalized to cell number, as measured by Quant-iT PicoGreen^™^ (Invitrogen).

### Statistical analysis

Major findings were replicated using a second cell line whenever possible. Data were expressed as mean ± SEM (standard error of the mean) of at least three independent experiments. Comparisons between groups were determined using an unpaired, two-tailed Student *t-*test (* *p* < 0.05; ** *p* < 0.01; *** *p* < 0.001 **** *p* < 0.0001). The one-way or two-way ANOVA test was used for comparisons between more than two groups. GraphPad Prism 9.2.3 software was used for statistical analyses.

## Results

### IDH1 expression in melanoma patients

Analysis of TCGA (The Cancer Genome Atlas) database revealed increased wt*IDH1* mRNA expression in tissues from primary and metastatic melanoma patients, as compared to normal skin. Kaplan-Meier analysis of IDH1 showed that higher mRNA expression of IDH1 in tumors is associated with poor overall survival in patients **(Fig. 1A-1C)**. Increased IDH1 expression was also observed at mRNA and protein levels in multiple human melanoma cell lines, as compared to normal melanocytes **(Fig. 1D and E)**.

**Figure 1:**
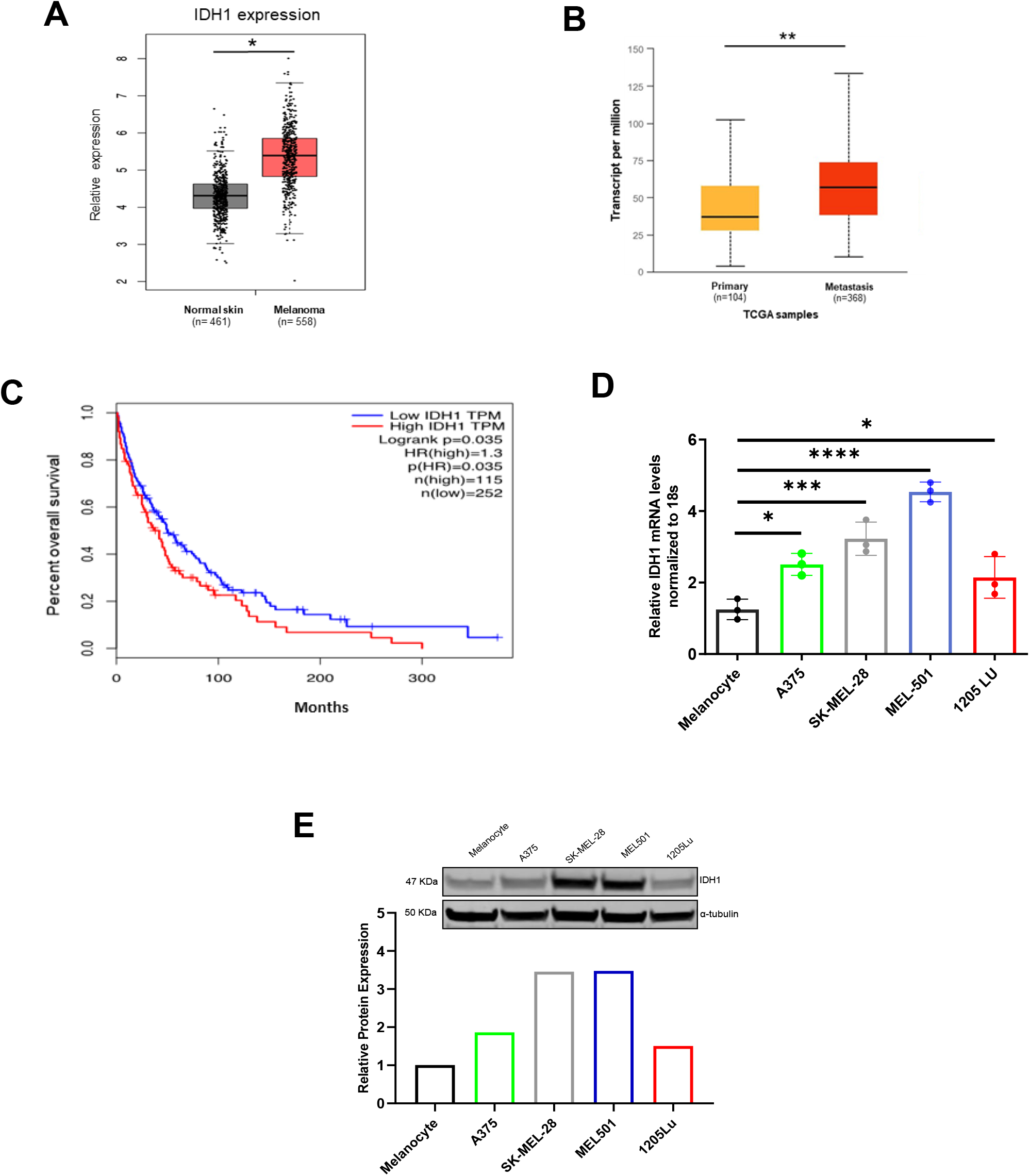
Wild-type IDH1 is overexpressed in primary and metastatic melanoma. **A]** RNA-sequencing data showing expression of IDH1 in human primary melanoma compared to that in normal skin tissues, *P < 0.05. **B]** IDH1 RNA expression in human primary melanoma compared to metastatic melanoma *P < 0.05. **C]** Correlation between IDH1 expression and overall survival rate of melanoma patients by Kaplan-Meier analysis using the log-rank test P<0.035. The data of Fig. A, B, and C were obtained from the TCGA database. **D]** *IDH1* mRNA expression level in different melanoma cell lines by qPCR. Expression levels are normalized to 18S expression in each cell line **E]** Immunoblot analysis of IDH1 in different melanoma cell lines and primary human melanocytes; alpha-tubulin used for normalization of cellular protein. The relative protein level of IDH1 is quantified by densitometry. Each data point represents the mean ± SEM of three independent experiments. *, P *<* 0.05; **, P *<* 0.01; ***, P *<* 0.001).

### IDH1 impacts growth and antioxidant defense under nutrient withdrawal in melanoma cells

Human melanoma cells were first confirmed to contain wtIDH1 genomic sequence (**Fig. 2A, Supplementary Fig. S1A and S6A)**. Cells were subsequently cultured in normal tissue culture media (25 mM glucose, supraphysiologic) or low glucose conditions (2.5 mM). Glucose withdrawal led to an acute increase in IDH1 mRNA and protein expression in melanoma cells **(Fig. 2B, Supplementary Fig. S1B)**, as an adaptive metabolic response that was previously observed in pancreatic cancer cells cultured under similar conditions (20). IDH1 siRNA silencing (**Fig. 2C Supplementary Fig. S1C**) led to more than a two-fold increase in ROS levels under low glucose conditions. The effect was negligible under high glucose conditions, highlighting the expendability of the enzyme under nutrient abundance (**Fig. 2D, Supplementary Fig. S1D**). 8-OHdG analyses revealed increased levels of DNA oxidation after siRNA suppression of IDH1, particularly under low glucose (**Fig. 2E** and **Supplementary Fig. S1E)**.

**Figure 2:**
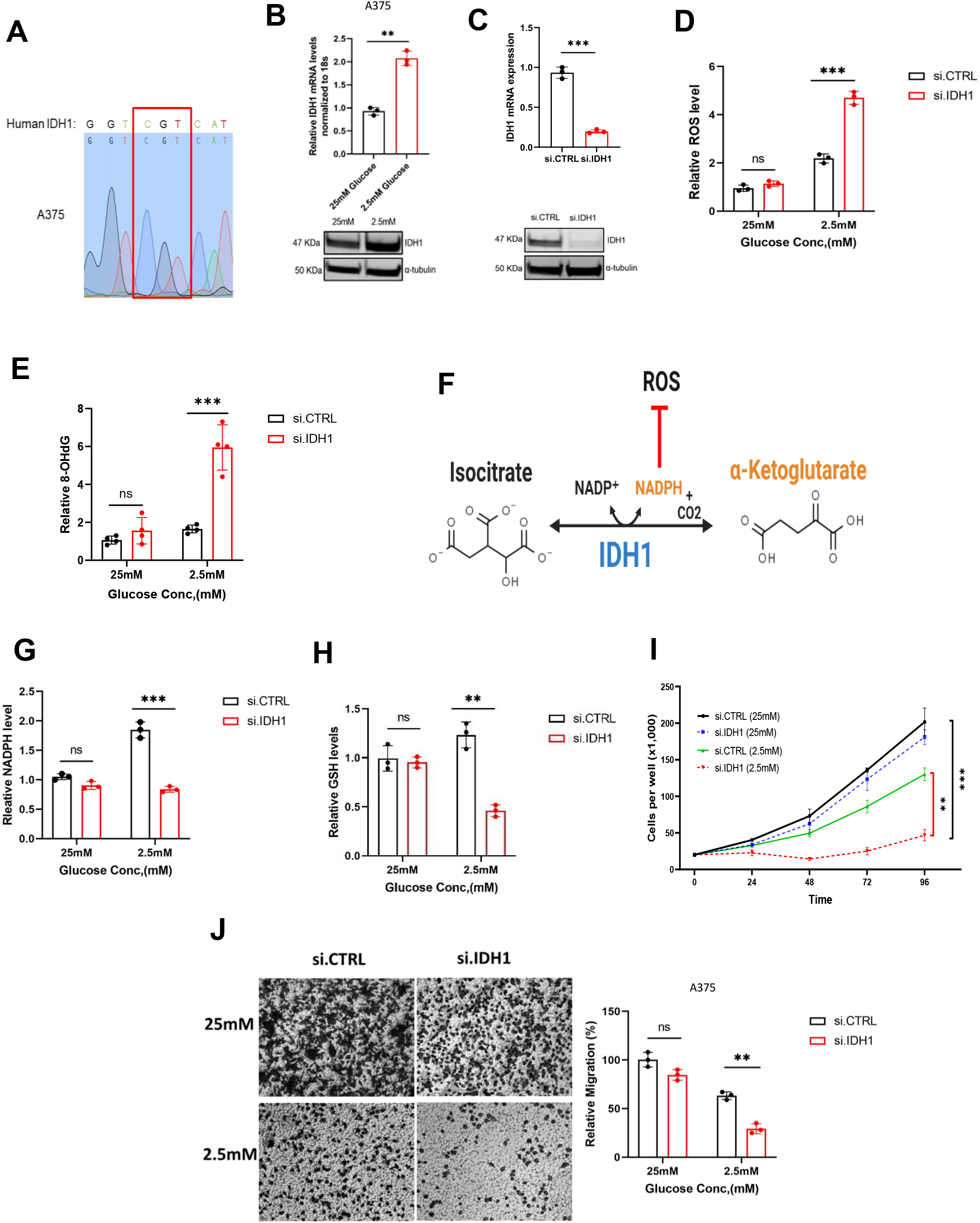
IDH1 knockdown suppresses melanoma cell growth and induces ROS under glucose withdrawal. **A]** Sanger sequencing of PCR amplicons correlated with codon 132 of the IDH1 gene in A375 cells. **B]** qPCR and immunoblot analysis for IDH1 expression in A375 under 2.5 mM glucose compared with 25 mM glucose for 48 hours. **C]** qPCR and immunoblot analysis for IDH1 expression after IDH1 silencing by siRNA oligos (si.IDH1) compared with control (si.CTRL) in A375 cells. **D]** Relative ROS levels in si.CTRL and si.IDH1 A375 cells for 48 hours under the indicated glucose concentrations. **E]** Relative 8-OHdG levels in DNA extracted from A375 cells under indicated conditions for 48 hours. **F]** Schematic of the IDH1 enzymatic reaction. **G]** Relative NADPH levels in A375 cells cultured under the indicated conditions for 72 hours. **H]** Relative GSH levels in si.IDH1 and si.CTRL A375 cells under the indicated glucose concentration. **I]** Cell viability (trypan blue assays) of A375 after silencing IDH1 compared to control (si.CTRL) under high and low glucose conditions for the indicated time points. **J]** Representative cell images of A375 (4X magnification) and quantification of transwell migration under the indicated conditions after silencing IDH1 compared to control. Each data point represents the mean ± SEM of three independent experiments. N.S., nonsignificant; *, P < 0.05; **, P < 0.01; ***, P < 0.001; ****, P < 0.0001.

The impact of IDH1 on cancer cell antioxidant defense was likely attributable to enhanced generation of NADPH related to upregulated IDH1 expression (**Fig. 2F**) observed with low glucose stress (**Fig. 2G**). IDH1 silencing abrogated the increase in NADPH (**Fig. 2G**). Similarly, siRNA suppression of IDH1 diminished GSH levels under low glucose conditions, but lacked impact under glucose abundance (**Fig. 2H**). Cell proliferation studies mirrored these results. Under low glucose conditions, IDH1-deficient cells failed to proliferate, yet IDH1-deficient melanoma cell growth was unaffected under high glucose conditions **(Fig. 2I Supplementary Fig. S1F)**. Along these lines, IDH1 expression similarly affected cell migration of melanoma cells under low glucose conditions (**Fig. 2J and Supplementary Fig. S1G)**.

### Metabolic changes associated with IDH1 expression

Under glucose withdrawal, parental melanoma cells experience increased mitochondrial respiration (OCR) and reduced glycolysis (ECAR) (**Supplementary Fig. S2A and S2B**), showing the importance of mitochondrial metabolism under nutrient scarcity. Suppression of IDH1 blocked this adaptive reprogramming and enhanced oxidative stress. Liquid chromatography coupled tandem-mass-spectrometry (LC-MS/MS) metabolomics in melanoma cells cultured under low glucose conditions revealed distinct metabolomic profiles **(Fig. 3A and Supplementary Fig. S2C)**. Hierarchical clustering and heatmap profiling of the top 50 altered metabolites **(Fig. 3B and 3C**) demonstrated reductions in mitochondrial TCA metabolites and associated metabolites of the TCA cycle (ATP and NADH), as well as a redox shift reflective of significant oxidative stress under low glucose. Of note, the two products of IDH1 oxidative decarboxylation (αKG and NADPH) were both reduced with IDH1 suppression, and upstream reactants (citrate, isocitrate and NADP+) were increased **(Fig. 3B and 3C)**. Pathway enrichment analysis revealed additional dysregulation of other metabolic pathways, including pyrimidine synthesis and glutamine/glutamate metabolism **(Fig. 3D**). Some of these changes were apparent under high glucose, but were less pronounced (**Supplementary Fig. S2D**). Consistent with these findings, siRNA against IDH1 under low glucose conditions substantially reduced OCR in melanoma cells **(Fig. 3E and 3F)**, with negligible effects under high glucose conditions. **(Supplementary Fig. S2E**).

**Figure 3:**
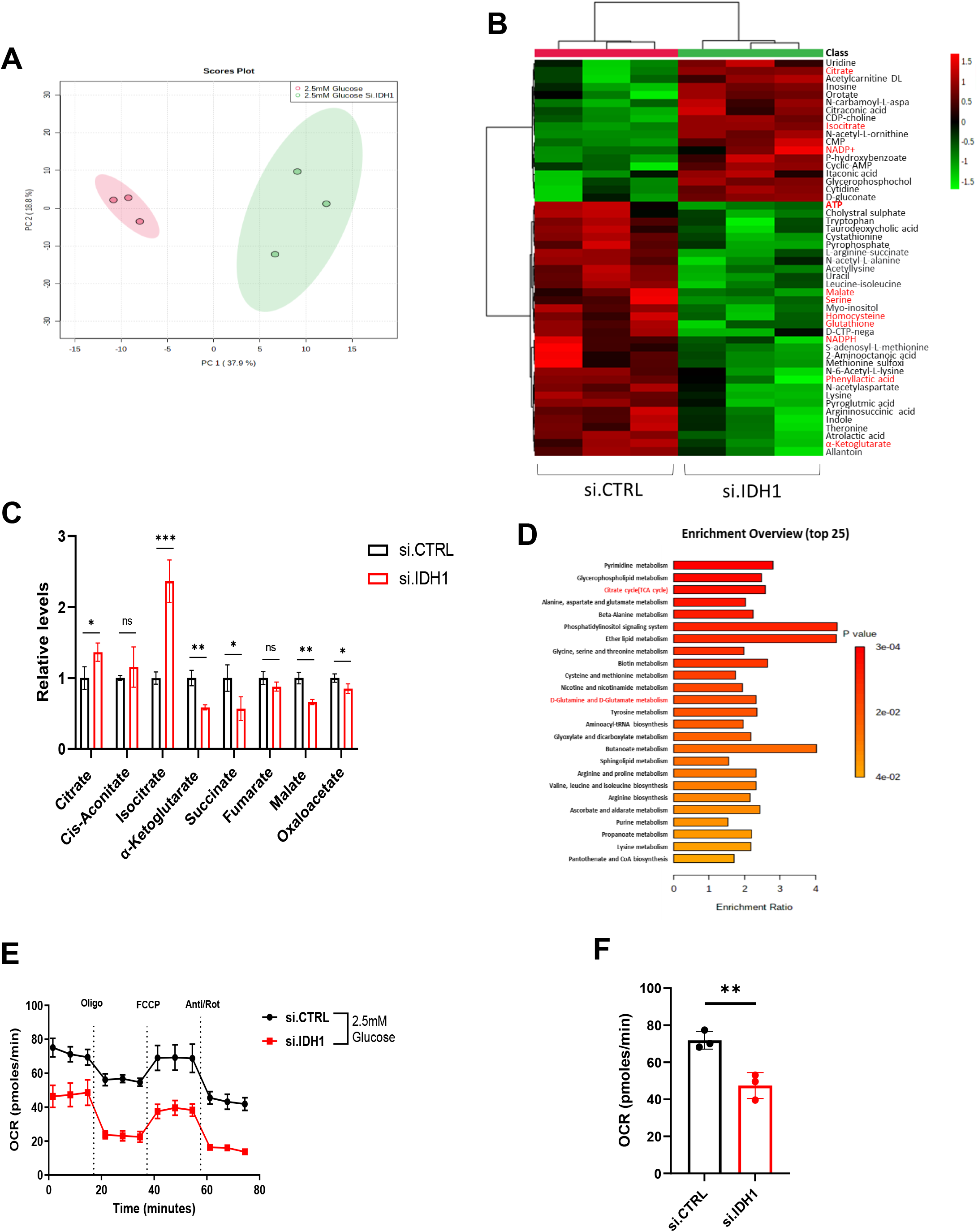
IDH1 supports mitochondrial function under stress. **A]** Principal-component analysis (PCA) of metabolites analyzed by LC-MS/MS performed on A375 cells, after transfection with si.IDH1 and si.CTRL (n = 3 samples). **B]** A heatmap of the top 50 metabolites with the greatest change in A375 cells after transfection with si.IDH1 versus si.CTRL (n = 3 independent samples) under 2.5 mM glucose and analyzed by LC/MS. The scale is log 2 fold-change. **C]** Relative levels of TCA cycle metabolites from A375 after transfection with si.IDH1 and si.CTRL under 2.5 mM glucose for 12 hours. **D]** Metabolite set enrichment analysis of A375 cells. **E]** Representative oxygen consumption rate (OCR) in A375 cells transfected with si.IDH1 and si.CTRL, and cultured in 2.5 mM glucose for 24 hours. Treatment with mitochondrial inhibitors are indicated: oligomycin (Oligo), FCCP, antimycin A and rotenone (Anti/Rot) and **F]** Basal mitochondrial OCR. Each data point represents the mean ± SEM of three independent experiments. N.S., nonsignificant; *, P < 0.05; **, P < 0.01; ***, P < 0.001.

### Pharmacologic inhibition of IDH1 reduces cell viability and inhibits tumor progression

Ivosidenib (AG-120) is an FDA-approved drug developed to selectively target mutant IDH1 (39–41). We recently discovered that the drug potently inhibits wtIDH1 under low Mg^2+^ conditions in pancreatic cancer models, as the cation competes with the inhibitor for negatively charged amino acid residues at the allosteric site (24). Herein, we observed that AG-120 had minimal impact on melanoma survival in a clonogenic assay under high Mg^2+^ conditions in two separate cell lines. However, under low Mg^2+^ conditions, pharmacologic wtIDH1 inhibition paralleled the above results observed with IDH1 gene suppression. Treatment impaired melanoma survival at low glucose levels, since cells are dependent on wtIDH1 for antioxidant defense and mitochondrial function under this condition (**Fig. 4A and 4B; Supplementary Fig. S3A and S3B**). These results were recapitulated in an independent cell proliferation assay (**Fig. 4C and 4D**; **Supplementary Fig. S3C and S3D**).

**Figure 4:**
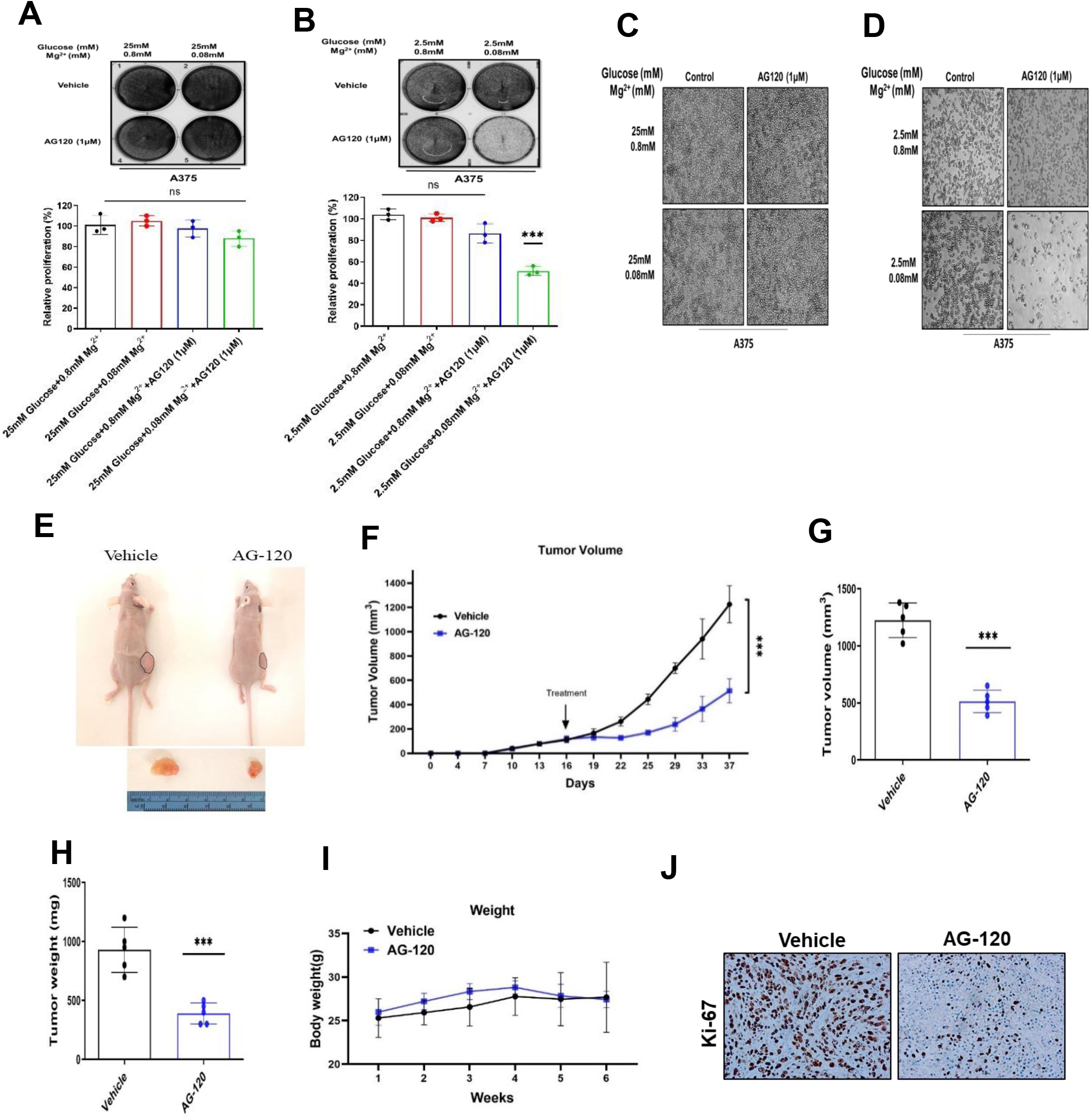
AG-120 is a potent wild-type IDH1 inhibitor under low glucose and magnesium conditions. A] Representative images of high glucose (25 mM) and **B]** low glucose (2.5 mM) colony formation assays in the A375 cell line. Cells were treated with vehicle control and AG-120 (1μM) under the indicated conditions, and stained with crystal violet solution. Quantification (%) is shown in the graphs. **C]** Under high glucose and **D]** low glucose conditions, phase-contrast images (4X magnification) were taken after treated with vehicle control or AG-120 (1μM) for 4 days under indicated nutrient conditions. **E]** Representative image of excised and in vivo tumors of A375. **F]** Tumor growth curves of A375 melanoma xenografts in nude mice. Tumor sizes were assessed twice per week using calipers (n=5 per group). **G]** Average tumor volume of A375 xenografts at the end of the experiment (day 26) (n=5 tumors per group). **H]** Average tumors weights (mg) of A375 xenografts in each group (n=5 per group). **I]** Body weights of A375 melanoma xenografts in nude mice (n=5 per group). **J]** Cell mitoses in tumor xenografts were estimated by nuclear immunolabeling (Ki-67). Scale bar, 50 μm. Each data point represents the mean ± SEM of three independent experiments. N.S., nonsignificant; *, P < 0.05; **, P < 0.01; ***, P < 0.001; ****, P < 0.0001.

IDH1 was subsequently deleted from melanoma cell lines using CRISPR/Cas9 editing **(Supplementary Fig. S4A, IDH1-KO)**. Equal numbers of IDH1-KO and control cells were injected into the flanks of nude mice. Unlike control cells, IDH1-KO cells failed to grow extensively *in vivo* (**Supplementary Fig. S4B-S4D**). Findings were replicated in pharmacologic studies. AG-120 was administered at the same dose previously used in animal studies of mutant IDH1 tumors (150 mg/kg orally twice a day) (42,43). Treatment significantly impaired tumor growth without any appreciable weight loss in the mice (**Fig. 4E-I**). Notably, intra-tumoral glucose and Mg^2+^ levels were markedly reduced in this model compared to adjacent normal skin and serum (**Supplementary Fig. S4E-S4F)**. Diminished cancer cell proliferation was validated by Ki-67 immunolabeling of harvested tumors (**Fig. 4J**).

### Targeting IDH1 sensitizes melanoma cells to chemotherapy

TMZ is one of the two most commonly used chemotherapeutics (the other being DTIC) in patients with advanced melanoma. In prior clinical studies, TMZ achieved a dismal objective response rate of just 14% (44). Since IDH1 inhibition enhances both total cellular ROS and oxidative damage within the nucleus (**Fig. 2D and E, Supplementary Fig. S1B and S1C**), we hypothesized that IDH1 inhibition would synergize with melanoma associated-chemotherapy known to exhibit oxidative and cytotoxic effects (45). We further hypothesized that the combination may be effective irrespective of glucose levels where chemotherapy replaces low glucose as a primer for oxidative stress, and therefore requires intact wtIDH1 for melanoma cell survival. In fact, IDH1 siRNA silencing resulted in a three- and nine-fold increase in TMZ sensitivity under high and low glucose conditions, respectively (**Fig. 5A and 5B, Supplementary Fig. S5A and S5B**).

**Figure 5:**
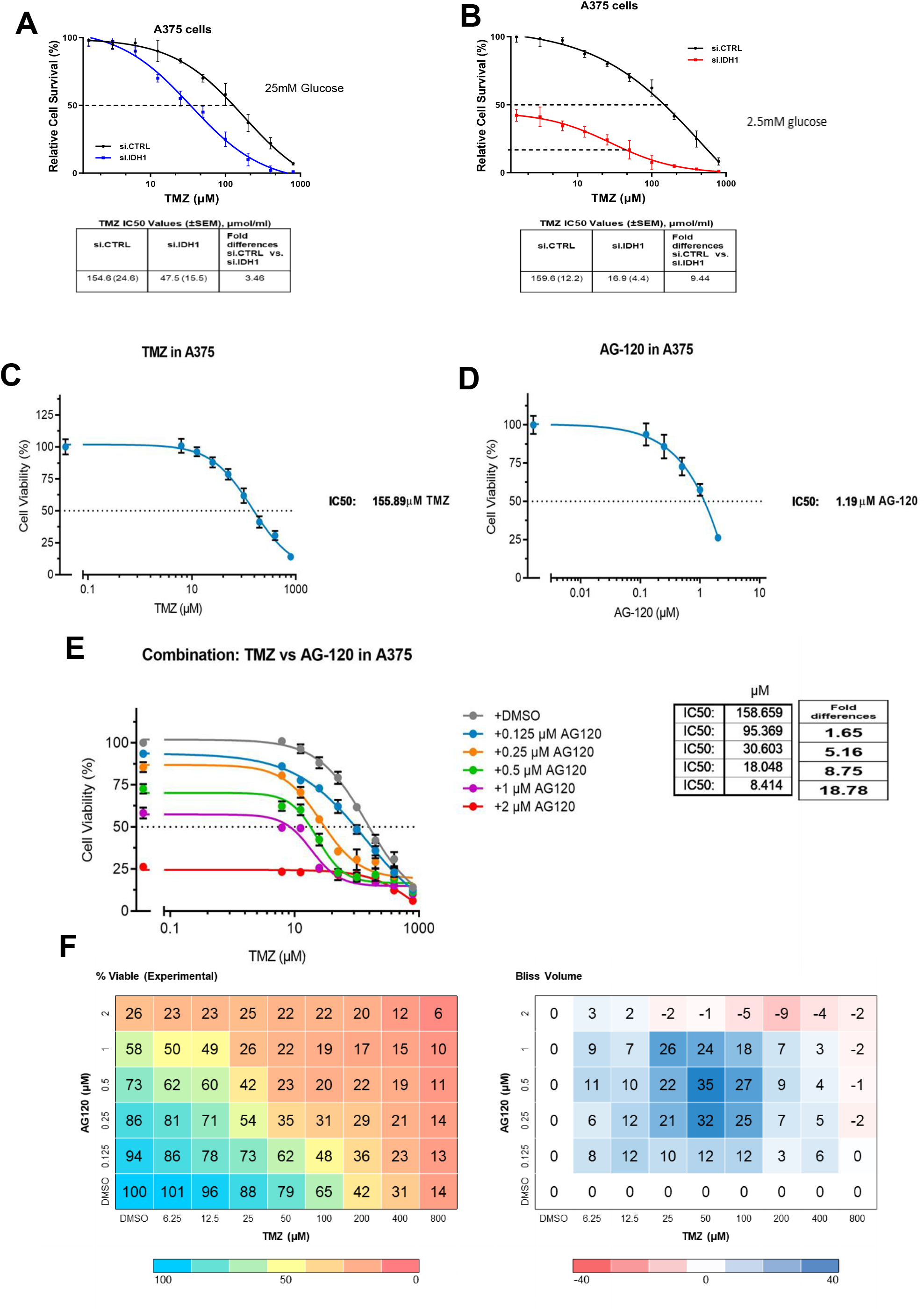
Targeting IDH1 sensitizes melanoma cells to conventional anti-melanoma cytotoxic therapy. **A]** Silencing IDH1 followed by treatment with TMZ for 5 days under high glucose (25 mM) and **B]** low glucose concentrations (2.5 mM) in the A375 cell line. IC_50_ values are provided. **C]** Cell viability of the A375 cell line treated with the indicated doses of TMZ. IC_50_ values are provided. **D]** Cell viability of the A375 cell line treated with indicated doses of AG-120. IC_50_ values are provided. **E]** Drug sensitivity in the A375 cell line under low glucose concentrations and with varying doses of TMZ and AG-120 cultured for 5 days. IC_50_ is provided. **F]** Drug matrix heatmap 5×8 (AG-120 and TMZ) grid showing percent viability and Bliss Independence scores in A375 cells cultured under 2.5 mM glucose for 5 days. Positive values reflect synergy and appear blue on the heatmap. All treatments with AG-120 were carried out under low glucose and low Mg^2+^ concentrations.

Pharmacologic IDH1 inhibition with AG-120 and TMZ phenocopied IDH1 silencing experiments. Targeting IDH1 in combination with chemotherapy treatment led to more than a three-fold increase in ROS levels under low glucose conditions (**Supplementary Fig. S5C and S5D)**. Dose response data using each drug alone **(Fig. 5C and 5D, Supplementary Fig. S5E and S5F)** informed the dosing used for eventual combination studies. These data revealed that pharmacologic IDH1 inhibition rendered TMZ substantially more potent (up to 18-fold at some dosing levels) in melanoma cell lines, A375 and SK-MEL-28 (**Fig. 5E and 5F, Supplementary Fig. S5G and S5H**). For instance, TMZ alone had an IC50 of 155.89 µM against A375 cells. The lack of potency is consistent with poor clinical efficacy. The addition of AG-120 at a dose slightly below its IC50 concentration (1 µM) shifted the TMZ IC50 downward by more than an order of magnitude (to 8.4 µM). As a result, a positive Bliss score with various dosing combinations was observed.

### IDH1 inhibition increases melanoma sensitivity to TMZ *in vivo*

The combination of these drugs given to mice also revealed enhanced anti-tumor activity *in vivo*. Mice bearing either human melanoma A375 cells transplanted into nude mice (**Fig. 6**) or murine B16-F10 melanoma cells into C57BL/6J mice (**Supplementary Fig. 6**) were treated with vehicle, TMZ (30 mg/kg intraperitoneal once a day), AG-120 (150 mg/kg orally twice a day), or combination AG-120+TMZ (150 mg/kg orally twice a day + 30 mg/kg intraperitoneal daily) **(Fig. 6A and Supplementary Fig S6B)**. While AG-120 was more effective than conventional chemotherapy as a single-agent, the combination was by far the most effective, as evidenced by both a reduction in tumor growth (**Fig. 6B-D)** and survival **(Supplementary Fig. 6C)**. The drug combination was well-tolerated by mice, without any reduction in body weight. (**Fig. 6E**). The effect of the combination was validated molecularly by a substantial reduction in Ki-67 immunolabeling in harvested tumors (**Fig. 6F**) and a dramatic increase in cleaved caspase-3 immunolabeling (**Fig. 6G**).

**Figure 6:**
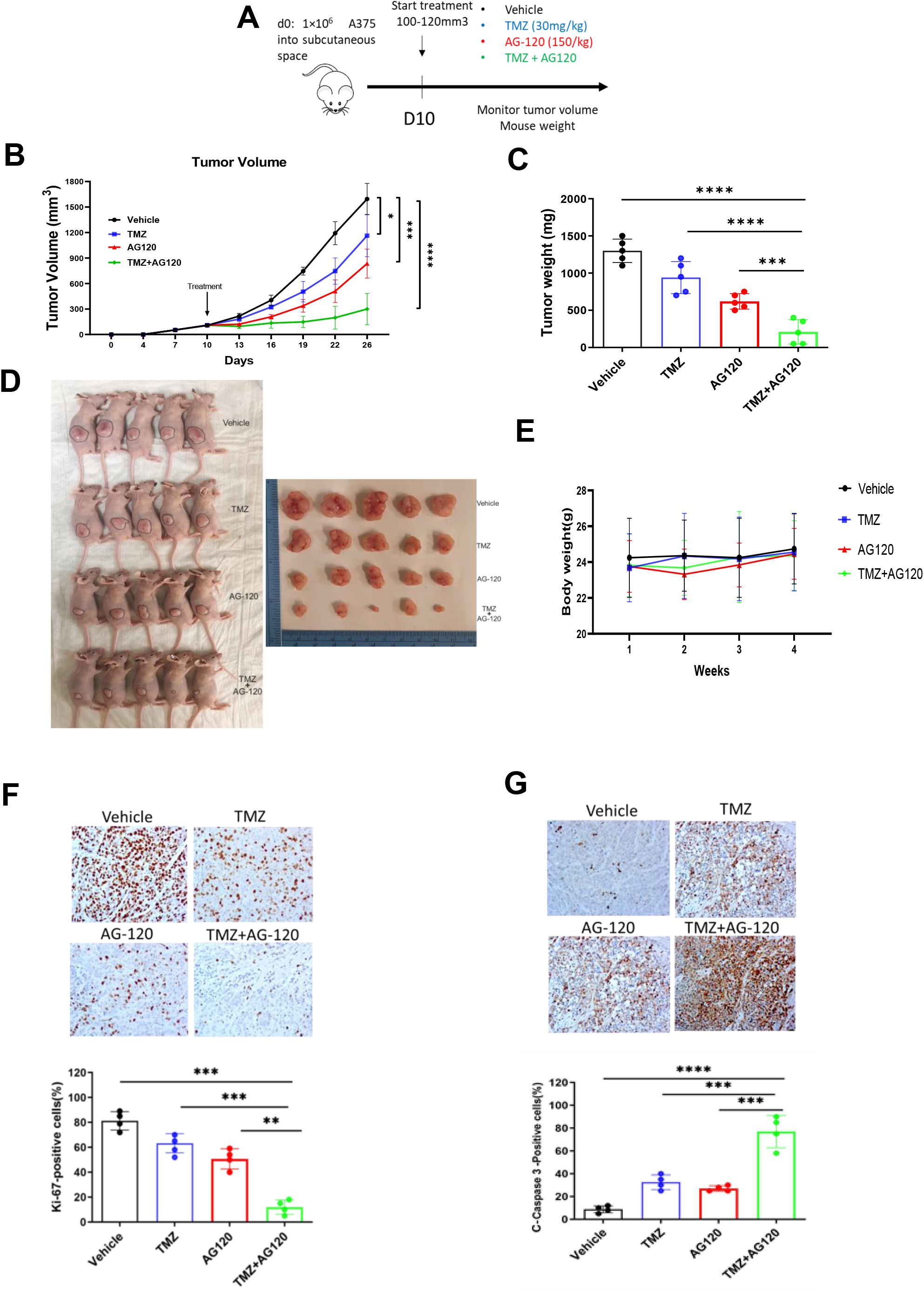
Treatment of mice bearing melanoma xenografts with TMZ in combination with AG-120. **A]** Schematic represents the treatment model after 1×10^6^ A375 melanoma cells were injected subcutaneously into the flank of nude mice. A separate experiment with B16-F10 melanoma murine cells involved 4×10^4^ cells injected subcutaneously into C57BL/6J recipient mice. After 8-10 days, when tumors reached 100-120 mm^3^, mice were divided into four groups and treated with i) Vehicle; ii) TMZ (30 mg/kg) every day; iii) AG-120 (150 mg/kg) twice a day; iv) AG-120 +TMZ (150 mg/kg + 30 mg/kg). **B]** Tumor growth curves of A375 melanoma xenografts in nude mice. Tumor sizes were assessed twice per week using calipers (n=5 per group), **C]** Average tumors weights (mg) of A375 xenografts in each group (n=5 per group) **D]** Representative image of excised and in vivo tumors of A375. **E]** Body weights of A375 melanoma xenografts in nude mice (n=5 per group). **F]** Cell proliferation in tumor xenografts was estimated by nuclear immunolabeling (Ki-67). Scale bar, 50 μm. Quantitation is shown below from four random fields per section. **G]** Tumor xenograft apoptosis was estimated with labeled cleaved caspase-3. Quantitation is shown below from four random fields per section. Scale bar, 50 μm. Each data point represents the mean ± SEM of at least three independent experiments. *, P < 0.05; **, P < 0.01; ***, P < 0.001; ****, P < 0.0001.

## Discussion

Tumor cells grow in harsh metabolic microenvironments and require robust molecular strategies to combat oxidative stress for survival (14). It follows then that interventions supporting the neutralization of reactive oxygen species would favor tumor growth. In fact, multiple studies in mice and humans reveal that antioxidant supplementation increases tumor growth and promotes progression in diverse cancer types, including melanoma, suggesting a rescue effect (46–49). This observation can be leveraged for therapeutic purposes. Targeting key components of antioxidant defense in tumors would conversely thwart cancer progression and metastasis by exacerbating the threat imposed by oxidative stress (38,39).

We show here that wtIDH1 functions towards this end and represents a metabolic vulnerability. Indeed, a small number of compelling studies previously cast a light on the enzyme as a promising therapeutic target in cancer. The pioneering work in this group of studies actually employed *in vitro* melanoma models but was never further pursued until the present work. Metallo et al. first observed that under hypoxic conditions, wtIDH1 activity was critical for melanoma cell survival because it encouraged the IDH1 reaction to favor a reductive direction (to the left in **Fig. 2F**) (23). That is, in the absence of glucose withdrawal, αKG derived from glutamine was converted into isocitrate and propelled carbon substrate towards *de novo* lipogenesis and tumor growth. Targeting IDH1 with siRNAs impaired cell proliferation under those conditions. Later, Jiang et al. showed that reductive carboxylation of glutamine was also important for anchorage-independence in tumor spheroids of different cancer types, and that this was again highly dependent on wtIDH1(21). Isotope tracer studies suggested that the cytosolic isocitrate produced by wtIDH1 through the reductive reaction was transferred to mitochondria, where oxidation back to αKG by mitochondrial IDH2 augmented NADPH and minimized mitochondrial ROS. Calvert et al. was the first to demonstrate that wtIDH1 favors oxidative decarboxylation (to the right in **Fig. 2F**) in certain tumor models (e.g., glioblastoma) to produce cytosolic NADPH for antioxidant defense and ROS control (52). Targeting IDH1 augmented oxidative stress and reduced glioblastoma growth.

We recently validated the importance of wtIDH1 in diverse pancreatic cancer models, and established several key principals in that work. First, wtIDH1 was especially important for cancer cell survival under nutrient limiting conditions. Both NADPH and αKG produced by the oxidative decarboxylation of isocitrate (to the right in **Fig. 2F**) were critical for adaptive survival under these conditions. Second, both oxidative IDH1 products mechanistically support mitochondrial function, in addition to antioxidant defense, and this wtIDH1 function was also essential for cancer cell survival under metabolic stress. αKG serves a key anaplerotic role in support of TCA cycling and mitochondrial function, while NADPH reduces mitochondrial ROS. Third, AG-120 and other allosteric wtIDH1 inhibitors developed to selectively target the mutant IDH1 isoenzyme, are actually potent wtIDH1 inhibitors in tumors due to two specific conditions common to the tumor microenvironment: low Mg^2+^ levels which permit stronger binding of the compounds within the allosteric site of wtIDH1, and low nutrient levels (e.g., glucose in particular) which increase cancer cell reliance on the wild-type isoenzyme. The presence of these specific conditions render cancer cells vulnerable to allosteric IDH1 inhibition.

In the present study, we sought to build on the prior work to more firmly establish wtIDH1 as a therapeutic target in melanoma and leverage novel insights to propose an immediately translatable therapeutic strategy for patients. In this study, wtIDH1 appeared to be important for melanoma survival under nutrient limited conditions, and genetic ablation of the enzyme slowed tumor growth in mouse melanoma models. Further, findings observed with IDH1 suppression were phenocopied by AG-120 treatment.

Cytotoxic chemotherapeutics remain the treatment backbone across most tumor types (53– 57), yet have been largely abandoned for melanoma. Currently, chemotherapy usage is limited to patients with metastatic melanoma after disease progression or drug intolerance with immunotherapy or oncogene-targeted agents (58). Cytotoxic treatment options include TMZ (alkylating agent), DITC (alkylating agent), paclitaxel (or albumin-bound paclitaxel) (microtubule inhibitor), and carboplatin (platinum agent and DNA cross-linker) (59). The most commonly used among these agents, TMZ (an oral prodrug of DITC) and DITC, have roughly equivalent activity against melanoma (60). Progression-free and overall survivals associated with these agents in patients with advanced melanoma are dismal and frankly unacceptable-just 2 and 7 months, respectively. Less than 20% of patients survive beyond 2 years without the benefit of newer therapies (2). These results have prompted investigations into mechanisms to induce chemosensitivity in melanoma as a strategy to offer readily available second or third-line options for patients with refractory disease. A common thread among many of the studies is the use of adjuvants that promote oxidative stress as a mechanism to reduce chemotherapy resistance in melanoma cells (30,61). In this study, IDH1 inhibition with AG-120 potently induced oxidative stress in melanoma cells under nutrient limitation (**Supplementary Fig. S5C and S5D)**, and effectively synergized with conventional anti-melanoma chemotherapy in cell culture and in mouse melanoma models.

In conclusion, IDH1 inhibition profoundly impairs the growth of melanoma cells in culture and xenografts in mice. IDH1 suppression enhances ROS and impairs mitochondrial function in tumors. As a result, wtIDH1 inhibition with AG-120 was effective against melanoma tumors, especially in combination with a conventional anti-melanoma cytotoxic agent (TMZ). Future studies aimed at validating these findings in additional melanoma cancer models will provide an even stronger rationale to test AG-120 (ivosidenib) and chemotherapy in patients with refractory metastatic melanoma.

## Authors’ Contributions

**M. Zarei:** Investigation, data curation methodology, writing-original draft, writing–review and editing. **O. Hajihassani:** Investigation, methodology. **J. J. Hue:** Investigation, methodology, **H. J. Graor:** Investigation, methodology. **M. Rathore:** Investigation, methodology **A. Vaziri-Gohar:** Investigation, methodology. **J.M. Asara:** Investigation, methodology. **J.M. Winter:** Conceptualization, funding acquisition, supervision, administration, writing and editing. **L. D. Rothermel:** Conceptualization, funding acquisition, supervision, administration, writing and editing.

## Abbreviations

IDH1: Isocitrate dehydrogenase 1
TMZ: Temozolomide
ROS: Reactive oxygen species
TCGA: The Cancer Genome Atlas
TCA: Tricarboxylic acid cycle; α-
KG: alpha-Ketoglutarate
GSH: glutathione.

## Supplementary Information

**Supplemental Figure 1:**
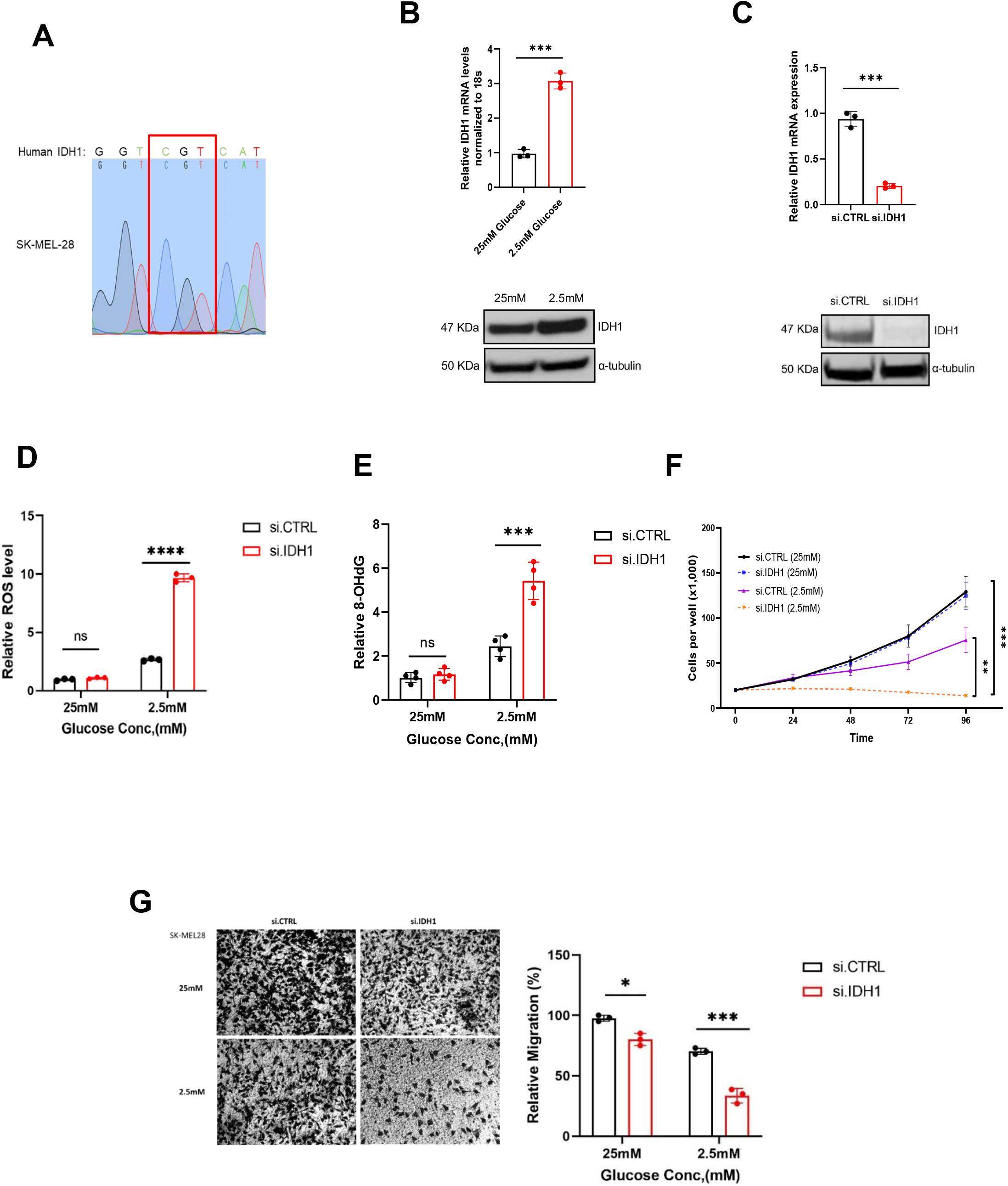
Validation that IDH1 suppression disrupts redox homeostasis and cell proliferation under glucose limitation in melanoma cells. **A]** Sanger sequencing of PCR amplicons correlated with codon 132 of the IDH1 gene in SK-MEL-28 cells. **B]** qPCR and immunoblot analysis for IDH1 expression in SKMEL-28 under 2.5 mM glucose compared with 25 mM glucose for 48 hours. **C]** qPCR and immunoblot analysis for IDH1 expression after IDH1 silencing by siRNA oligos (si.IDH1) compared with control (si.CTRL) in SK-MEL-28 cells. **D]** Relative ROS levels measured by DCFDA in SK-MEL-28 cells transfected with si.IDH1 and si.CTRL at the indicated glucose concentrations for 48 hours. **E]** Relative 8-OHdG levels in DNA extracted from SK-MEL-28 cells under indicated conditions for 48 hours. **F]** Cell viability (trypan blue assays) of SK-MEL-28 after silencing IDH1 compared to control (si.CTRL) under high and low glucose conditions for the indicated time points. **G]** SK-MEL-28 cell images (4X magnification) and quantitation of transwell migration under the indicated glucose concentrations.

**Supplemental Figure 2:**
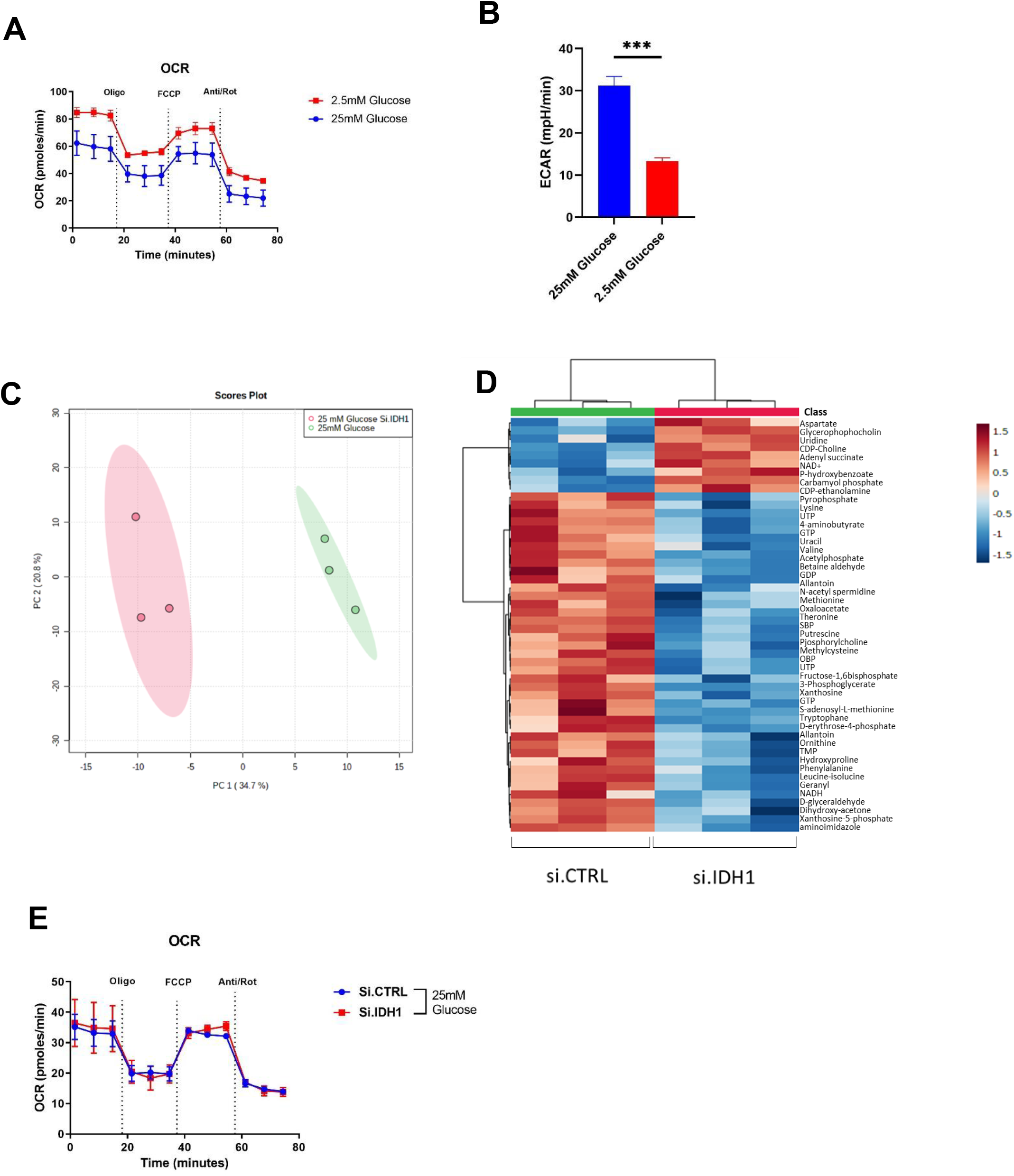
IDH1 supports mitochondrial function under glucose limitation. **A]** Representative OCR tracing in A375 melanoma cells cultured under the indicated glucose concentrations for 24 hours and **B]** Extracellular acidification rate (ECAR) response of A375 cells under the indicated conditions.**C]** PCA of metabolites analyzed by LC-MS/MS performed on A375 cells under high (25 mM) and low (2.5 mM) glucose (n = 3 samples). **D]** A heatmap of the top 50 metabolites with the greatest changes in A375 cells (n = 3 independent samples) under 25 mM glucose. The scale is log 2 fold-change. **E]** Representative OCR tracing in A375 melanoma cells cultured under the indicated glucose concentrations for 24 hours.

**Supplemental Figure 3:**
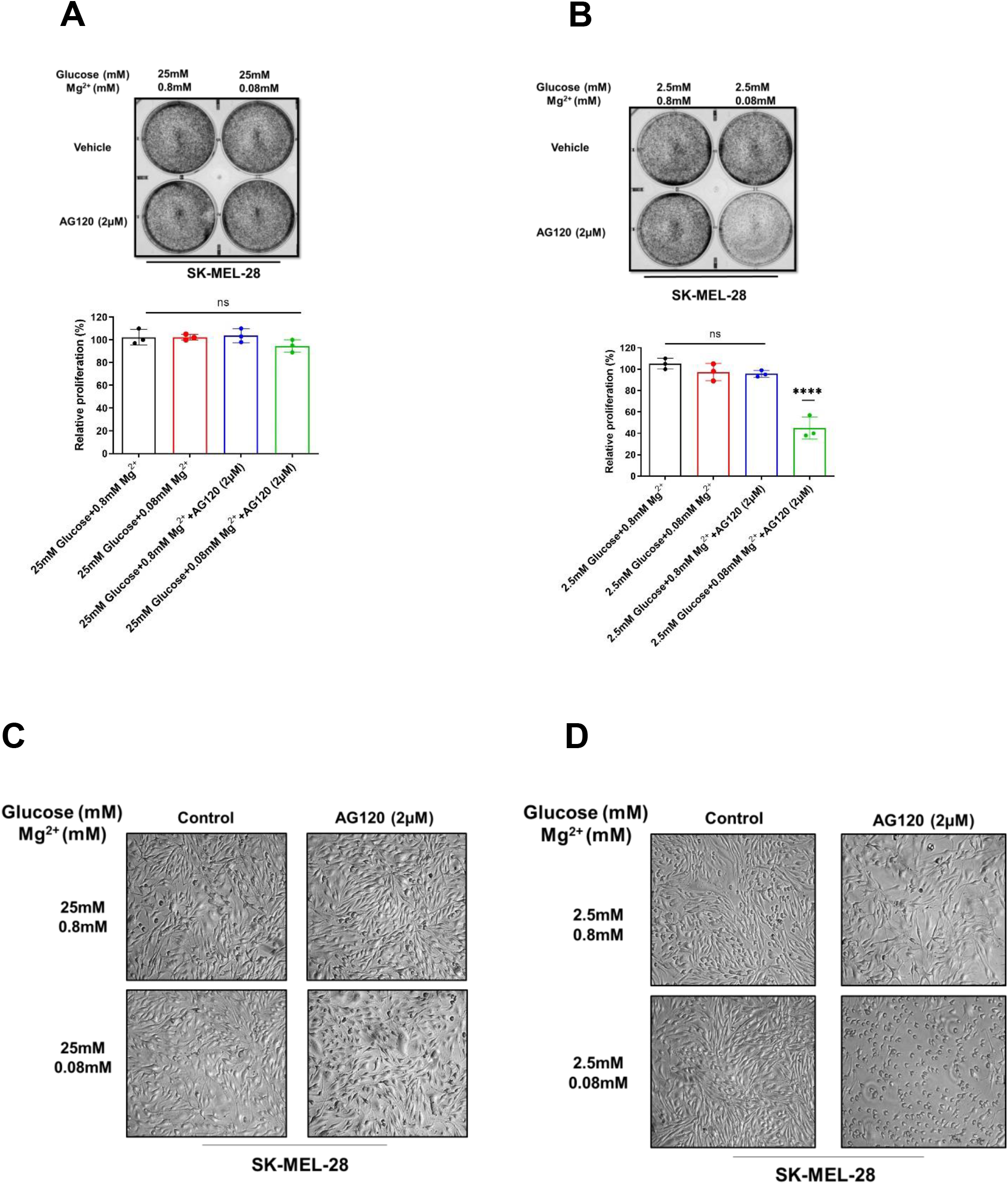
AG-120 is a potential wtIDH1 inhibitor under glucose limitation in SK-MEL-28 cells. **A]** Representative images of colony formation assays for cells cultured under high (25 mM) and **B]** low glucose (2.5 mM). The cells were treated with vehicle control or AG-120 (2 μM) under the indicated conditions for 9 days. Quantitation (%) is shown in the graph at the bottom. **C]** Cells cultured under high glucose, or **D]** low glucose were captured by phase-contrast imaging (4X magnification), after treatment with vehicle control or AG-120 (2 μM) for 4 days. Each data point represents the mean ± SEM of three independent experiments. N.S., nonsignificant; ****, P < 0.0001.

**Supplemental Figure 4:**
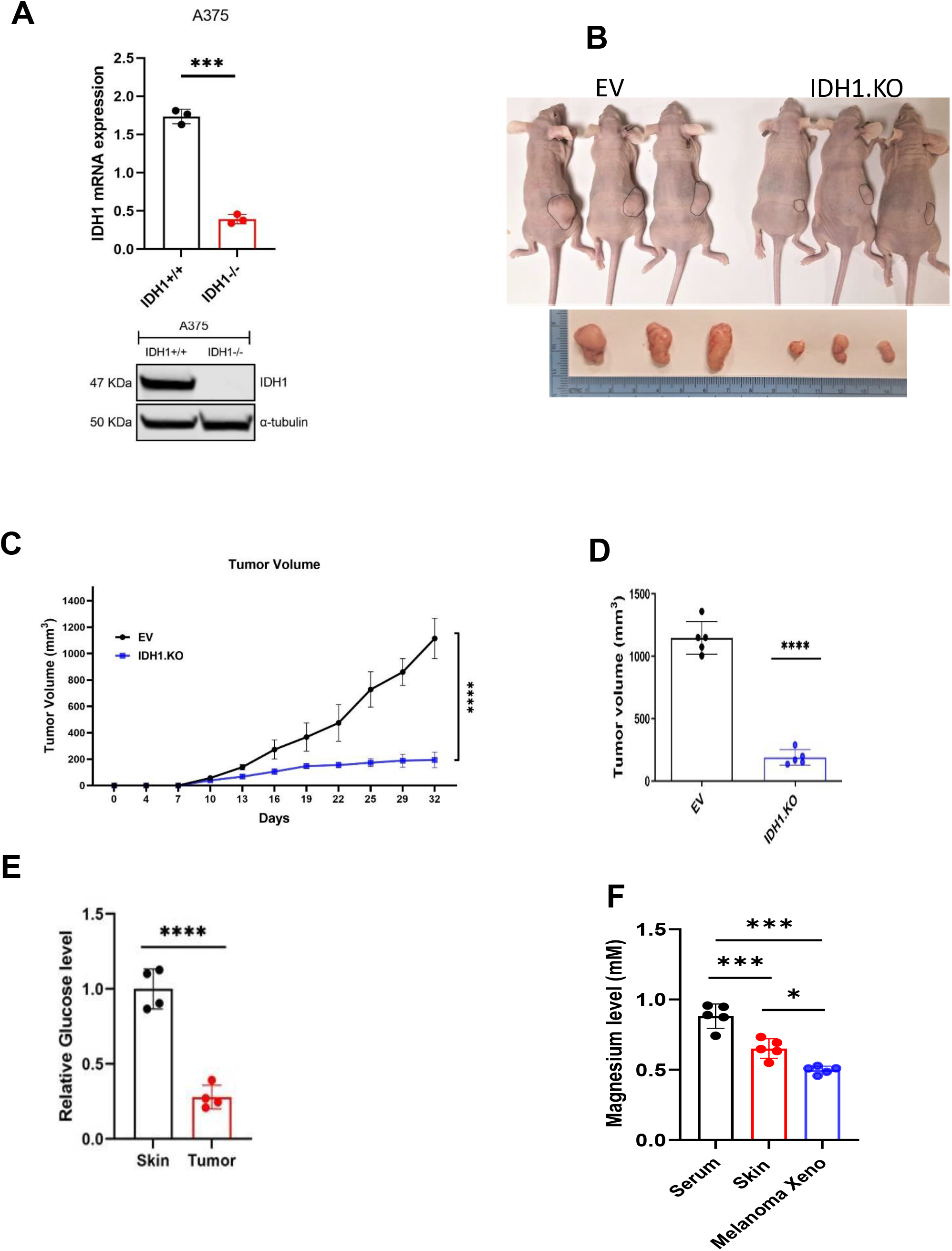
IDH1 knockout suppresses tumor growth *in vivo*. **A]** Relative mRNA levels, normalized to mRNA levels of 18S; Western blot analysis of IDH1 expression after IDH1 knockout by CRISPR/Cas9 (IDH1.KO) compared to control (IDH1.EV) A375 cells. **B]** Mice were injected with IDH1.EV and IDH1.KO A375 cells (n=5 per group) and tumor sizes were monitored for 5 weeks. Images of tumors at the end of the experiment are shown. **C]** Tumor volumes of IDH1.EV and IDH1.KO A375 xenografts. **D]** Histograms show tumor volumes with IDH1.EV and IDH1.KO A375 xenografts at the end of the experiment. **E]** Relative glucose levels in adjacent skin and xenograft. **F]** Relative free magnesium levels in skin, xenografts and serum. Each data point represents the mean ± SEM. *, P < 0.05; **, P < 0.01; ***, P < 0.001; ****, P < 0.0001.

**Supplemental Figure 5:**
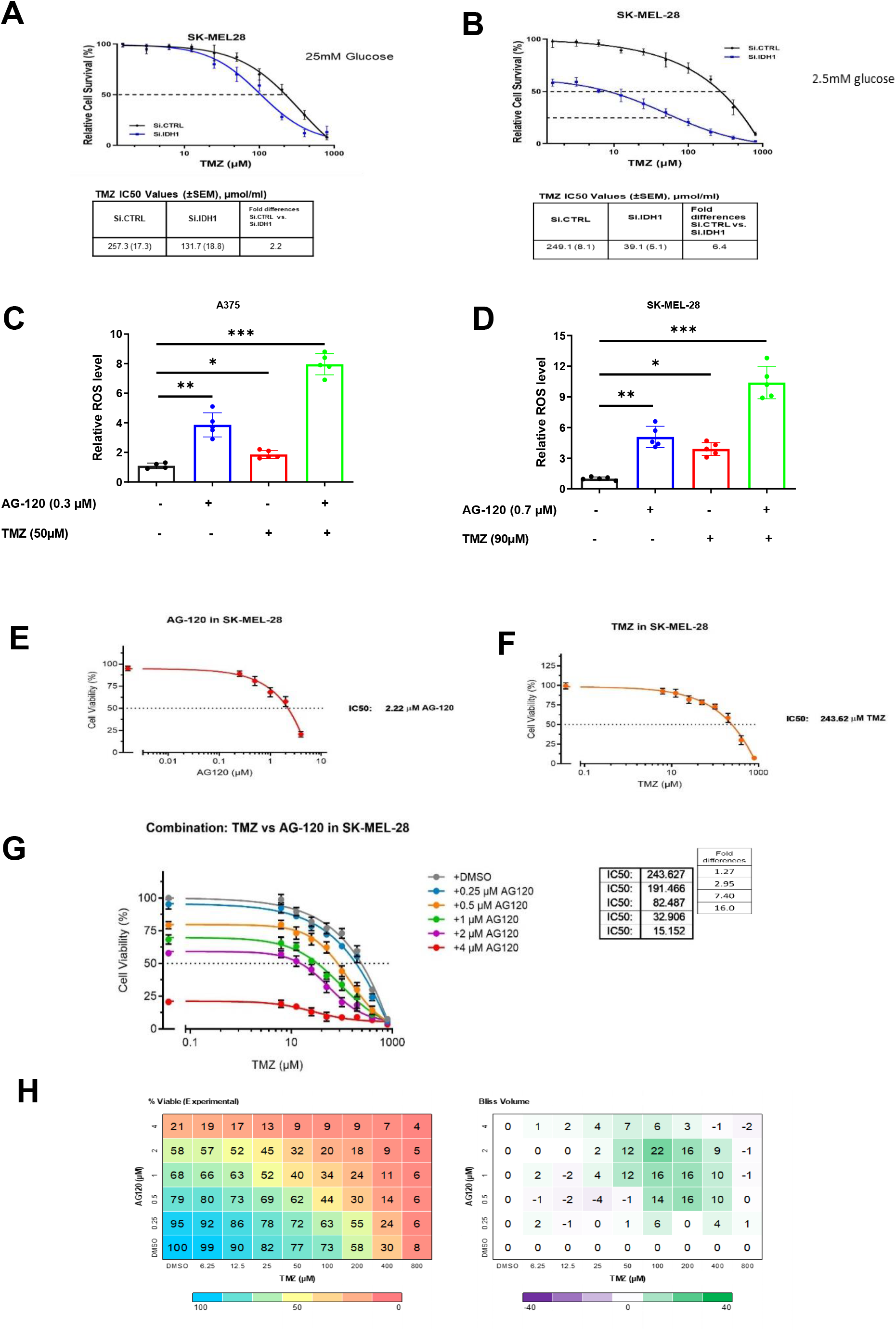
*In vitro* response of melanoma cells to treatment with TMZ in combination with AG-120. **A]** Silencing *IDH1* followed by treatment with TMZ for 5 days under high glucose (25 mM) and **B]** low glucose (2.5 mM) in SK-MEL-28 cells; IC_50_ values are provided. **C]** Relative ROS levels after 48 hours in A375 cell line under low glucose combined with TMZ and AG-120. **D]** Relative ROS levels after 48 hours in SK-MEL-28 cells under low glucose combined with TMZ and AG-120. **E]** Cell viability of SK-MEL-28 cells, treated with the indicated doses of TMZ. IC_50_ values are provided. **F]** Cell viability of SK-MEL-28 cells treated with indicated doses of AG-120 for 6 days. IC_50_ values are provided. **G]** Drug sensitivity in SK-MEL-28 cells under low glucose, with varying concentrations of TMZ and AG-120, cultured for 5 days under low glucose. IC_50_ results are provided. **H]** Drug matrix heatmap 5×8 (AG-120 and TMZ) grid showing percent viability and Bliss Independence scores in SK-MEL-28 cells cultured under 2.5 mM glucose for 5 days. Positive values reflecting synergism appear green on the heatmap (Bliss volume ≥ 10). All treatments with AG-120 were carried under low glucose (2.5 mM) and low Mg^2+^ (0.08 mM).

**Supplemental Figure 6:**
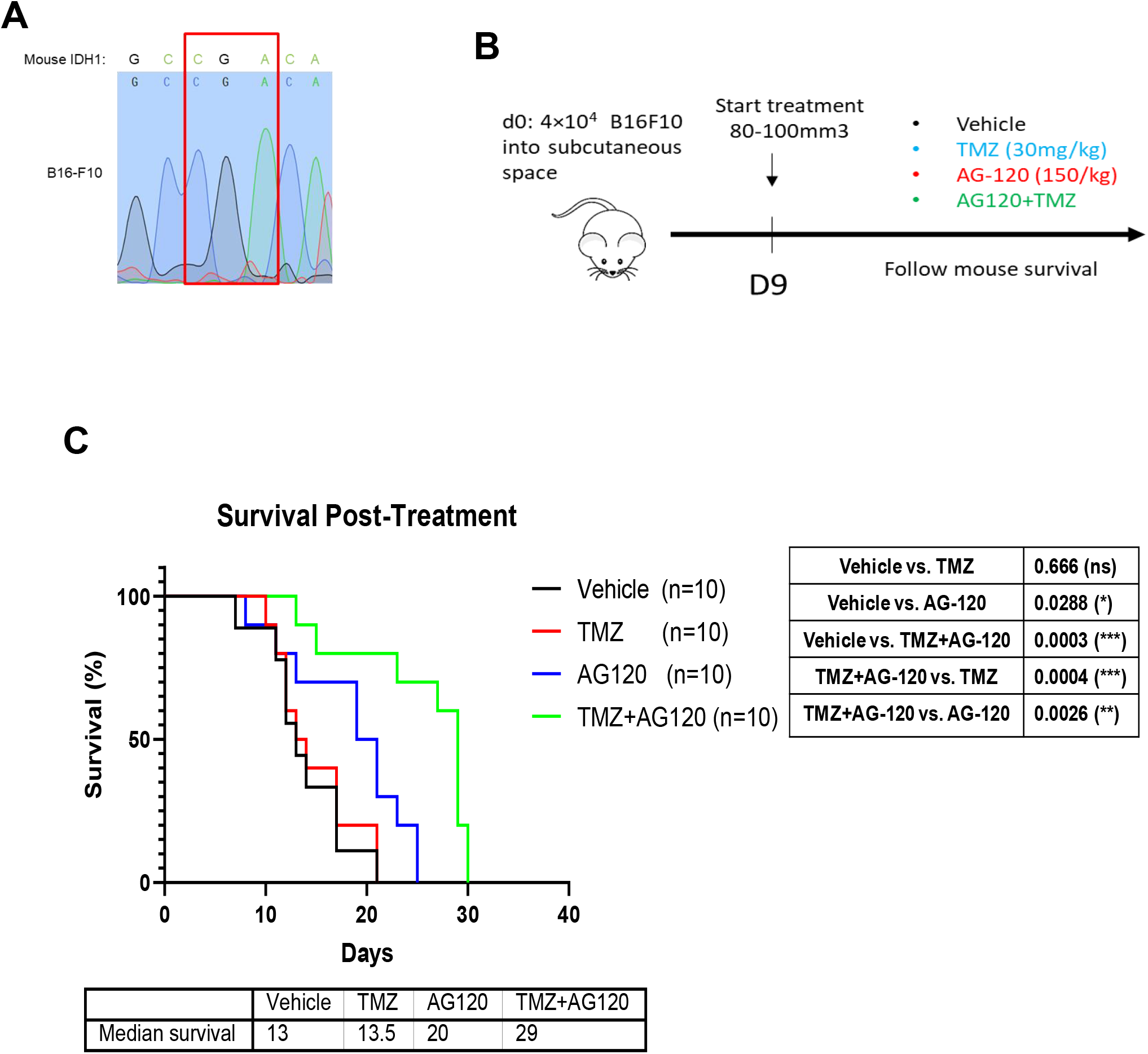
Treatment of mice bearing B16-F10 tumors with TMZ in combination with AG-120. **A]** Sanger sequencing of PCR amplicons correlated with codon 132 of the IDH1 gene in B16-F10 murine melanoma cells. **B]** Schematic represents the treatment model after 4×10^4^ B16-F10 melanoma murine cells were injected subcutaneously into the flanks of C57BL/6J recipient mice. After 9 days, when tumors reached 80-100 mm3, mice were divided into four groups and treated with i) Vehicle; ii) TMZ (30 mg/kg intraperitoneal once a day); iii) AG-120 (150 mg/kg orally twice a day); and iv) AG-120 + TMZ (150 mg/kg orally twice a day +30 mg/kg intraperitoneal daily). **C]** Survival data of C57BL/6J mice are represented by Kaplan-Meier curves. Significance between each group was determined using the log-rank test.

## Graphical abstract

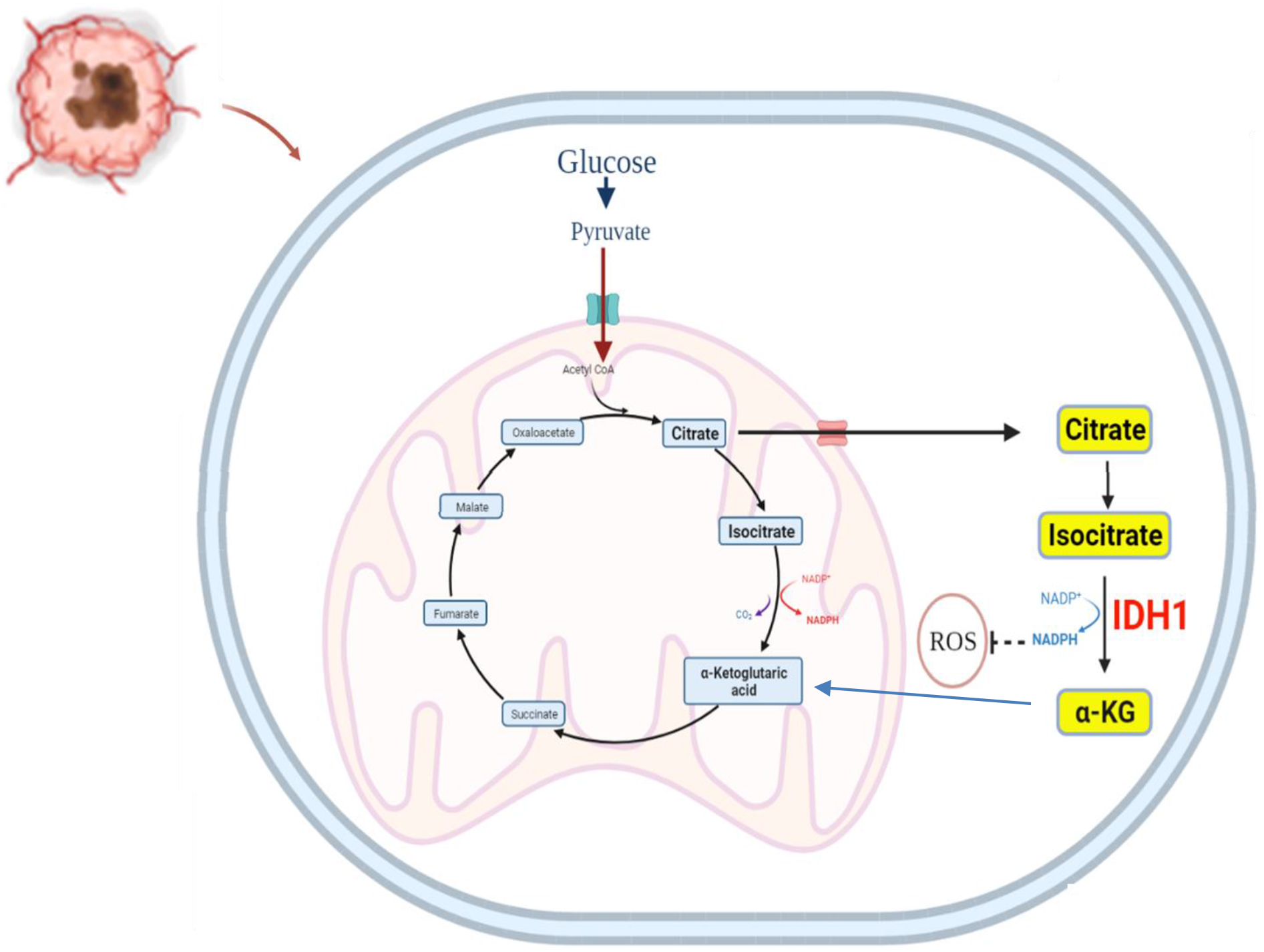
Schematic shows how wild type IDH1 expression and activity change under metabolic stress to enhance reductive power and mitochondrial function.

